# Fast dose fractionation using ultra-short laser accelerated proton pulses can increase cancer cell mortality, which relies on functional PARP1 protein

**DOI:** 10.1101/571703

**Authors:** E. Bayart, A. Flacco, O. Delmas, L. Pommarel, D. Levy, M. Cavallone, F. Megnin-Chanet, E. Deutsch, V. Malka

## Abstract

Radiotherapy is a cornerstone of cancer management. The improvement of spatial dose distribution in the tumor volume by minimizing the dose deposited in the healthy tissues have been a major concern during the last decades. Temporal aspects of dose deposition are yet to be investigated. Laser-plasma-based particle accelerators are able to emit pulsed-proton beams at extremely high peak dose rates (~10^9^ Gy/s) during several nanoseconds. The impact of such dose rates on resistant glioblastoma cell lines, SF763 and U87-MG, was compared to conventionally accelerated protons and X-rays. No difference was observed in DNA double-strand breaks generation and cells killing. The variation of the repetition rate of the proton bunches produced an oscillation of the radio-induced cell susceptibility in HCT116 cells, which appeared to be related to the presence of the PARP1 protein and an efficient parylation process. Interestingly, when laser-driven proton bunches were applied at 0.5 Hz, survival of the radioresistant HCT116 p53^−/−^ cells equaled that of its radiosensitive counterpart, HCT116 WT, which was also similar to cells treated with the PARP1 inhibitor Olaparib. Altogether, these results suggest that the application modality of ultrashort bunches of particles could provide a great therapeutic potential in radiotherapy.

## Introduction

Among radiotherapy modalities, the use of hadrons (i.e. protons and heavier ions) for malignant tumor treatment takes advantage from their energy transfer characteristics in comparison to widely used X-rays. Hadron therapy allows high conformation of the dose distribution to the target volume because of the sharp lateral penumbra and distal dose fall-off, as most of the energy is deposited within the Bragg peak. These properties make it possible to increase dose deposited in the tumor target while reducing dose in adjacent healthy tissues, preventing associated sides effects^1, 2^.

During the past decade, important goals have been reached on laser-driven proton sources, making them a reliable alternative for *in-vitro* studies. The biggest difference between laser-driven sources and conventional ones is the temporal structure of the irradiation. While conventional proton sources deliver a continuous beam at a dose-rate of several Gy/min, laser-driven particle beams are delivered as separate ultra-short bunches, typically in the range of nanoseconds, and dose rates as high as 10^9^ Gy/s^3–5^. Laser sources at Hz repetition rates are hence capable of delivering comparable average dose rates, whereas peak dose rate is 6 to 9 orders of magnitude higher.

While development of laser-driven proton sources is still ongoing, to reach energies relevant for clinical applications, it is crucial to characterize the radiobiological effects of pulsed ionizing radiation at high dose rate. Although the biological effects of proton irradiation on living systems have been widely studied^6^, much still has to be explored on the impact of protons delivered in such short pulses of ultra-high dose rates on living cells or tissues. During the last decade, several experimental campaigns proved the feasibility of radiobiological studies on intense laser facilities and were able to evaluate the biological effectiveness of such beam^3–5, 7–13^. These studies suggest that the radiobiological effectiveness of laser-driven protons is roughly similar to conventional beams, when considering DNA damaging potential or tumor cell killing. We recently described a set-up of four permanent magnet quadrupoles to shape and control the proton beam generated by the multi-terawatt laser SAPHIR at LOA, the only (French) laser-plasma infrastructure dedicated to radiobiology studies, and validated the robustness of the system by irradiating radiosensitive colorectal cancer cell line^5^. Here we confirmed the efficiency of laser-plasma protons beams compared to conventional ones on radioresistant glioblastoma cell lines, for which proton therapy is indicated. As a further step, we investigated the biological impact of the temporal aspect of laser-driven proton bunches. Despite the challenging implementation of radiobiology assays on a laser facility, we show that the variation of proton bunch repetition rate is associated with an oscillation of cell survival, which is found to be dependent on the PARP1 (poly ADP-Ribose polymerase 1) protein activity in tumor cells. This is the very first time the temporal structure of laser beam, that we called fast dose fractionation, is investigated and demonstrated to provide an increased therapeutic potential.

## Results

### Laser driven protons (LDP) are as efficient as conventional accelerated protons (CAP) and X-rays in inducing DNA double strand breaks and cell killing on glioblastoma cell lines

The favorable ballistic of proton beams make such treatment efficient for brain, base-of-skull and head-and-neck tumors. As previous experiments were performed on rodent, HeLa, lung or colorectal cells^3–5, 7, 8, 10, 11^, we decided to study the impact of LDP on the highly resistant glioblastoma cells lines, SF763 and U87-MG, with regard to CAP or X-rays. We first compared LDP-induced DNA double strand breaks (DSBs). DSBs were detected by microscopy through immunodetection of the histone H2AX phosphorylation on Ser139 (γH2AX). Cells were fixed one or 24 hours after three and six LDP’s bunches (corresponding to 2.1 ± 0.42 and 4.2 ± 0.84 Gy respectively, see methods section) or two and four Gy of CAP or X-rays (Fig.1A). As expected the amount of DSBs increased as the dose increased; it decreases between 1h and 24h, which reflects the DNA repair process. Similar amount of foci were quantified for LDP, CAP or X-rays one hour after irradiation either in SF763 or U87-MG cells (Fig.1B and 1C). Twenty four hours post-irradiation, no differences were observed for SF763 while the number of residual foci seemed to be higher for LDP compared to CAP and X-rays in U87-MG cells. However, this apparent difference was not significant. These results suggest that LDP induce DNA double strand breaks in resistant glioblastoma cell lines as well as CAP or X-rays.

To follow-up on the radiobiological consequences of LDP on cancer cells, we evaluated next how cell survival was affected by LDP. For that purpose, the two glioblastoma cell lines were irradiated with 0 to 12 proton bunches (corresponding to doses ranging from 0 up to 10 Gy) of LDP and the resulting dose response survival curves were compared to those obtained with cells irradiated with CAP or X-rays. It can easily be noticed that, for the SF763 cells, the survival curves corresponding to each irradiation condition overlap (Fig. 1D), giving similar D_10_ (doses giving 10% of cell survival, p values were > 0.3, table 1); the same result was observed with the U87-MG cell line. These data indicate that the impact of LDP on cell survival is similar to the one of CAP or X-rays. These results show that no significant difference in DNA double strand breaks detection and cell survival measurement could be detected between LDP, CAP and X-rays in highly resistant glioblastoma cell lines. These observations are in agreement with previous studies using others cell lines^3–5, 7, 8, 10, 11^ and confirm that laser-driven protons are able to affect cancer cells, at the global DNA damage and cell survival levels, with the same effectiveness compared to conventional accelerated particles.

**Table 1.** Comparison of doses giving 10% of cell survival (D_10_ values) from LDP, CAP and X-rays. D_10_ mean values ± SEM extracted from curves obtained in figure 1D and 1E are reported. Each value represents the mean of at least three independent experiments. Comparison using by two way ANOVA multiple comparisons test (Tukey’s multiple comparisons test) gave, at least, p>0.33.

| Radiation | SF763 | U87-MG |
| --- | --- | --- |
| LDP | 7.45±0.31 | 7.47±0.32 |
| CAP | 7.90±0.18 | 7.11±0.16 |
| X-rays | 7.95±0.24 | 7.45±0.06 |

**Figure 1.**
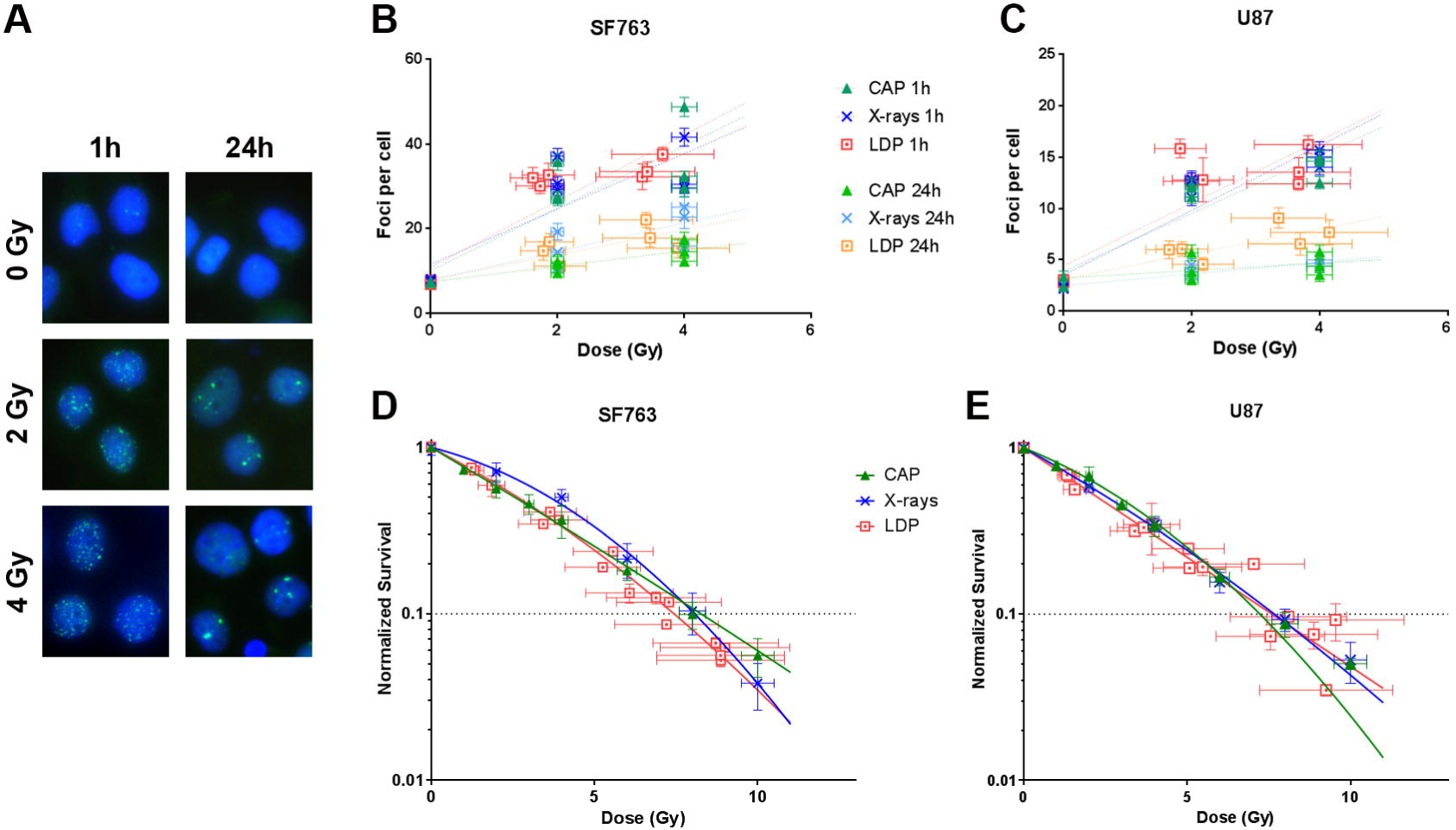
Dose responses of DNA damage foci formation and of cell survival. (A) Representative immune-fluorescent images of SF763 cells obtained 1h and 24h after exposure to the indicated doses of laser driven protons (LDP, dotted square), conventional accelerated protons (CAP, triangles) and X-rays (x cross). The negative controls (0 Gy) were sham-irradiated. Merged images show γH2AX (green) and DNA (blue). (B) and (C) Number of γH2AX foci induced by LDP, conventional accelerated protons CAP and X-rays respectively in SF763 and U87-MG cells. Each data point represents the mean of at least three independent experiments in which at least 300 nuclei were analyzed and averaged. (D) and (E) Normalized cell survival resulting from exposure to increasing doses LDP, CAP and X-rays respectively in SF763 and U87-MG cells. Each data point represents the mean of three replicates obtained at least with three independent experiments. Survival curves were generated following the linear quadratic model (R>0.96 and R>0.97 for SF763 and U87-MG cell lines respectively).

### Changing the dose delivery modality by modifying the repetition rate of LDP’s bunches can result in cell survival oscillation

Laser-driven proton acceleration in our configuration (see methods) reaches a maximum repetition rate of 0.5 Hz, i.e. a minimum of 2 seconds between to consecutive proton bunches. This delay can be increased to any value with a resolution of 200 ms. In our experiment, the cell lines under test (glioblastoma SF763 and U87-MG, colorectal cancer cells HCT116-WT and HCT116-p53^−/−^) were exposed to a fixed number of LDP bunches (five or nine for colorectal cancer and glioblastoma cell lines respectively). Different samples were irradiated with a variable delay between shots, from 60 to 2 seconds. Hence the total irradiation time varied from 15 to 540 seconds. Cell survival of each cell lines was monitored and reported as a function of the delay between LDP’s bunches. As shown in figure 2, such variation did not affect the survival neither of SF763 cells (p>0.531, Fig. 2A) nor of U87-MG cells (p>0.222, Fig. 2B). In the case of HCT116 cells, variation of proton bunches cadency affected the cell survival. The same outcome was observed with both HCT116 WT and its radioresistant counterpart HCT116 p53^−/−^; as expected the survival of HCT116 p53^−/−^ cells remained higher than the WT. For the two cell lines however the observed surviving fraction was a non-monotonic function of the delay between each pulse, with the apparition of two different maxima (p≤0.0038 and p<0.0001 for HCT116 WT and HCT116 p53^−/−^ respectively). Interestingly, the cell survival difference between HCT116 p53^−/−^ and HCT116 WT was maximum when the curves reached their maxima, then the two converged together at the shortest delays (at 2s p=0.9982). This observation suggests that an increase of proton bunch cadency could lead, depending on cell type, to an increase in cell mortality.

**Figure 2.**
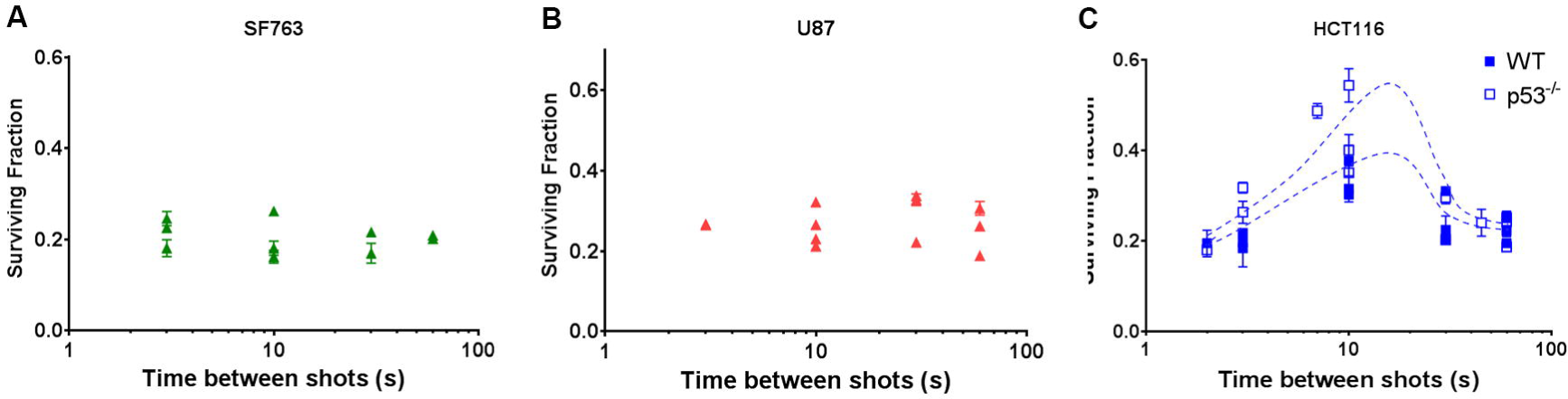
Cell survival dependency to the variation of the bunch repetition rate. Normalized survival fraction resulting from exposure for a given dose of LDP bunches delivered with different delay ranging from 3 to 60 seconds. Radioresistant Glioblastoma cell lines SF763 (triangles) (A) and U87-MG (dots) (B) were submitted to a dose corresponding to nine bunches of LDP (6.3±1.39 Gy). (C) radiosensitive colorectal cancer HCT116 cells, WT (white squares) and p53^−/−^ (black squares) were exposed to five bunches of LDP (3.5±0.77 Gy). Each data point represents the mean of three replicates obtained in at least three independent experiments.

### Non-monotonic character of cell survival as a function of LDP proton bunches cadency is related to the presence of functional PARP1 protein

About two decades ago, it was observed that cells exposed to two briefs pulses of relativistic electrons exhibited variable survival rates as a function of the delay between the two pulses. This phenomenon is called W-effect (WE) from the shape of the survival rate oscillation, where the delay was varied between few minutes and sub-second^14^. This process appeared to vary considerably among different cell lines and was shown to be dependent on an active PARP1 (poly(ADP-ribose) polymerase) protein^15^. The variation of the cell survival we observed in HCT116 cells by varying LDP proton bunch repetition rate could share origins with the W effect. However, this non-monotonic behavior was not observed in the other tested cell lines, which seems to indicate that this variation of cell susceptibility could also be linked to the PARP1 protein. Oxidative stress promotes the formation of large amounts of base damage leading to DNA breaks, which induces the over-activation of PARP1 and leads to protein parylation^16^. Thus, HCT116, SF763 and U87-MG cells were exposed to hydrogen peroxide (H_2_O_2_), then Western-Blot analyses were performed to detect the PARP1 protein and the related parylation. As shown in figure 3A, the PARP1 protein was detected in HCT116 WT cells as well as in HCT116 p53^−/−^ cells. When exposed to H_2_O_2_, parylation was induced, demonstrating the functional activity of PARP1 in these two cell lines. On the contrary, in U87-MG cells, very low amount of PARP1 could be detected and only extremely weak parylation was observed. Surprisingly, in SF763 cells, the PARP1 protein was efficiently detected, however, no increase of parylation was observed after cell exposition to hydrogen peroxide. This result suggests that in this cell line, although present, the PARP1 protein was not functional hence not able to promote parylation. The PARP1 protein activation and the related parylation process are crucial for DNA damage repair pathways activation^17^. Inhibition of PARP1 in cells where PARP1 is active produces an increased sensitivity to genotoxic stress, particularly in response to ionizing radiations^18^. To exploit this advantage for the treatment of cancer, PARP1 inhibitors, such as Olaparib have been developed^19^.

**Figure 3.**
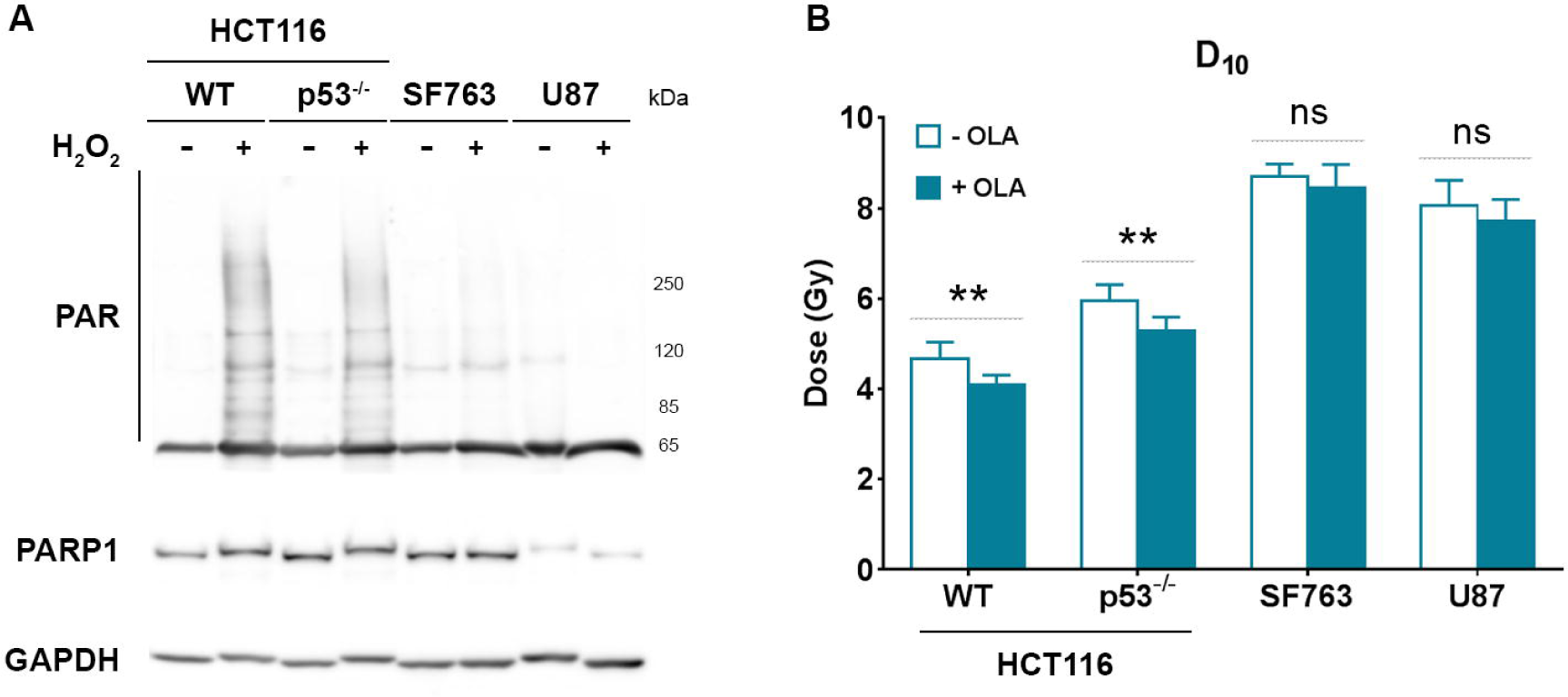
Determination of PARP1 status and related radiosensitization. (A) Western Blot detection of the PARP1 protein and parylation in total protein extracts from HCT116 WT, HCT116 p53^−/−^, SF763 and U87-MG cells left untreated or treated 10 min with 1mM hydrogen peroxide (H_2_O_2_). Protein amounts were normalized probing membrane with an anti-GAPDH antibody. (B) D_10_ values determined from survival curves obtained after exposure of HCT116 WT, HCT116 p53^−/−^, SF763 and U87-MG cells left untreated (white bars) or treated with 200 nM Olaparib (black bars) to increasing doses of X-rays. Survival curves were generated following the linear quadratic model (R>0.96); D_10_ mean values ± SEM are represented and were compared for each cell line using ratio paired t test. Each bar represents the mean of at least three independent experiments.

To further correlate the PARP1 activity status to the oscillating cell susceptibility to variation of the cadency of shots, cells from each of the cell lines previously used were treated or not with Olaparib (200nM) and then subjected to increasing dose of X-rays. The corresponding survival curves were generated and D_10_ values (doses giving 10% of cell survival) were extracted and compared (Fig.3B). As expected, in HCT116 WT cells, where PARP1 is functional, the use of Olaparib significantly increased cell mortality, which resulted in a decreased D_10_ values: from 4.68±0.36 to 4.09±0.22 (p=0.0066). The same outcome was observed treating HCT116 p53^−/−^ cells for which D_10_ value went from 5.95±0.36 down to 5.29±0.30 (p=0.0082). On the contrary, no significant variation of cell survival was detected with SF763 and U87-MG cells, exhibiting no or very few PARP1 activity, when submitted to the same treatment (D_10_=8.70±0.29 without and D_10_=8.45±0.52 with Olaparib treatment in SF763 cells, p=0.3558; D_10_=8.07±0.55 without and D_10_=7.72±0.48 with Olaparib treatment in U87-MG cells, p=0.1166). Together, these results suggest that the cell survival variation with the proton bunch cadency is related to the presence of a functional PARP1 protein.

### An increase of LDP’s bunches cadency can promote same cell radiosensitization as the use of PARP1 inhibitor

Cell survival level in response to variation of proton bunches repetition rate was correlated, in the different cell lines, to the presence of a functional PARP1 protein. It was verified next whether an inhibition of PARP1 would affected this variation or not. For this purpose, HCT116 WT cells were exposed, as in the previous experiment, to Olaparib (200nM) to inhibit PARP1, which led, as shown by western blot (fig.4A), to an absence of the H_2_O_2_-induced parylation. Cells were then subjected to five LDP’s bunches with delay ranging from 3 to 45 seconds.

**Figure 4.**
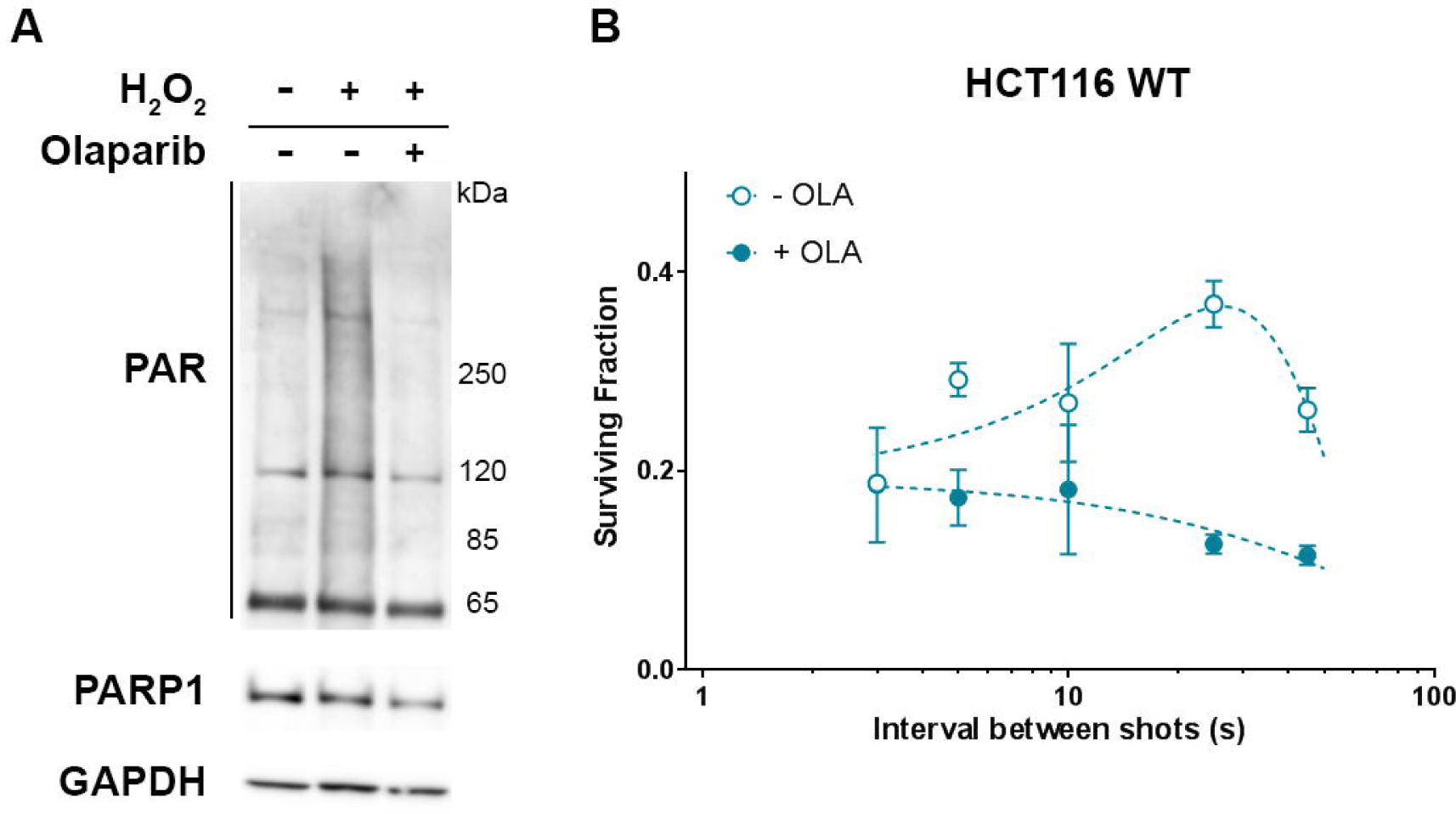
Impact of PARP1 inhibition on cell survival oscillation in response to the variation of the delay between LDP bunches. (A) Western Blot detection of the PARP1 protein and parylation in total protein extracts from HCT116 WT cells left untreated, treated 10 min with 1 mM hydrogen peroxide (H_2_O_2_) or treated one hour with 200 nM Olaparib before H_2_O_2_ treatment. (B) HCT116 WT cells were left untreated (black circles) or treated one hour with 200 nM Olaparib (white circles) and were then exposed to five bunches of LDP (3.5±0.77 Gy). Each data point represents the mean of three replicates obtained in two independent experiments. Comparisons of the cell survival were performed using by two way ANOVA multiple comparisons test (Tukey’s multiple comparisons test).

As expected, in absence of Olaparib, HCT116 WT cell survival variation was non-monotonic, increasing between 25 and 45 seconds to reach a maximum (p≤0.0316) before decreasing from 25 to 2 seconds where it reached a minimum. On the contrary, when cells were incubated in presence of Olaparib before exposition to LDP, cell survival variation was not detected any more (p≥0.0754). Interestingly, when bunches were separated by 2 seconds, the cell survival of cells only irradiated by LDP’s bunches reached the level obtained by those submitted to Olaparib treatment before LDP exposure, giving equivalent values (p>0,9999). This result first demonstrates that non-monotonic cell survival depends on the PARP1 protein activity. The PARP1 protein is a crucial actor in DNA damage signaling^20^, suggesting that cell survival variation induced by the LDP bunch repetition rate could be due to a specific impact of this dose deposition modality on DNA damage signaling and repair. Furthermore, we observed that an increase of the proton bunch cadency led cells exhibiting a functional PARP1 activity to the same decrease of cell survival level as combining LPD to PARP1 inhibition: this suggests that only using specific LDP dose deposition modality could be as efficient as drug-combined radiotherapy.

## Discussion

The main finding of this study is that more than the dose itself, the temporal dose deposition modality of LDP is a key feature, determining cancer cells response. Because of their favorable ballistic properties, hadrons (protons and carbon ions) represent a better alternative for the radiation therapy of solid cancer affecting organ-at-risk^1^. Due to the Bragg pick dose deposition profile, the dose is focused inside the target volume, reducing side effects to the neighboring healthy tissue. The development of laser plasma technology and its new paradigms for protons acceleration^21^ raises new possibilities for studying the effects of dose delivery modalities^22–24^.

Since the first proof of concept of irradiating cells with LDP^8, 12^, less than ten years ago, we and others have studied the biological impact of laser-accelerated protons and shown similar effectiveness of LDP, compare to conventional beams, to generate DSBs^3, 10^ and to induce cell killing^4, 5, 9, 11^. None of the previously existing studies, performed on various animal and human cell types, included highly resistant glioblastoma cells, one of the main indications of proton therapy. We observed, in agreement with the literature, similar DNA damaging potential and cell survival in response to LDP or conventional accelerators.

We and others have however reported a number of non-significant divergences which were considered to arise from differences in experimental procedures and endpoints (e.g. the dose applied), in reference photons and protons energies and LDP spectra. This study suggests that such discrepancies could also depend on the applied dose deposition modalities. When a sample is irradiated by laser-generated particles (electrons or protons), the total deposited dose is delivered as a sequence of multiple ultra-short bunches at ultra-high dose rates; the dose per bunch, hence the number of bunches required for a given dose, and the effective repetition rate depend on technical choices at the facility^3–5, 7–13^. Our study explores the radiobiological impact of the repetition rate of bunches and suggests that parameters such as the dose-per-bunch and the distribution in time of the LDP bunches should be better taken in account in future studies.

By varying the delay between proton bunches, we observed a non-monotonic variation of cell survival in HCT116 cells; this variation was not observed in SF763 nor in U87-MG cell lines. Similar variations with low energy electrons were already reported, two decades ago. Indeed, it has been shown, using a linear accelerator, that pulse irradiation brings about multiphasic, synchronous changes in the susceptibility of cells to a second pulse of radiation applied at variable intervals, lasting from fractions of second to a few minutes: these changes are at the origin of the W-shaped survival curve (W-effect) ^14, 25^. Further exploration of this phenomenon revealed that the WE depends on a functional PARP1 protein^15^. In our experiment, we also observed that the response of cell survival induced by the variation of the LDP bunches cadency was correlated to the presence of PARP1 and an efficient parylation process in cells exposed to H_2_O_2_. Hydrogen peroxide results in strong oxidative stress in cells, promotes the formation of large amount of base damage, leading to DNA breaks and protein parylation, which mostly accounts for PARP1^16^. Protein parylation in response to genotoxic stress is essential to promote DNA damage signalization and activation of DNA repair pathways^17, 26^. Consequently, the inactivation of the PARP1 results in enhanced genomic instability and increased radiation sensitivity^18^. This mechanism has been already exploited to develop drugs for cancer treatment. Several PARP inhibitors, such as Olaparib (AZD-2281, ASTRA-ZENECA), are currently tested in the clinic in combination with radiotherapy^27^. Consistently, in cells exhibiting normal parylation and PARP1 protein amount (HCT116 cells), the use of Olaparib combined with ionizing radiations led to an increased cell susceptibility that was not observed in SF763 and U87-MG, in which parylation was impaired. This existing parylation default explains why the U87-MG cell line was previously reported to be a non-responder cell line to PARP inhibitor^28, 29^. These results are consistent with the very recently published results showing that the deletion of PARP1 protein leads to resistance to PARP inhibitors^30^. They also explain why no difference of cell survival was observed for those cells when irradiations were performed with continuous beams. We then showed that, as for the WE, the bunches cadency related cell survival variation was abolished in response to PARP1 inhibition using Olaparib. This observation supports the idea that those two phenomena are related to DNA damage signaling or repair pathways. Indeed, it has been suggested that irradiation of cells at critical times of DNA damage recognition or repair following the first irradiation exposure may promote chromosome damage and long-lasting genomic instability^14, 15^.

The W-effect was demonstrated by irradiating cells with electron bunches at a dose rate of 12 Gy/s^14, 15^. Our observation, consistent with W-Effect and obtained with proton bunches, suggests this phenomenon to be likely related to the temporal feature of the dose delivery rather than the particles. The W-effect was observed after the irradiation with only two electron bunches. In our experimental conditions, excepted for the lowest dose of dose-response survival curves, the application of several pulses was required to reach the desire total dose. In previously published results with laser accelerators, dose was deposited either in one shot^3, 4^ or applying tens of shots depending on the available dose per shot of the infrastructure used^9–13^. As variation of cell survival appears to be correlated to multiple bunch irradiation, no differences could be observed with regards to conventional irradiation when cells were irradiated using a single bunch. In the case of dozen of bunches, the dose per bunch was very low ranging from 0.03 to 0.3 Gy per shot^9–13^. W-effect has been described applying shots of 1, 2 or 4 Gy^14, 15^ and, in our study, the SAPHIR parameters were set up to deliver a dose per bunch in the range of 1 Gy. These observations suggested that the variation of cell susceptibility in response to multiple bunch irradiation occurs only if each single shot bunch has a minimal dose close to the Gray or more, leading sufficient DNA damage and PARP1 activation.

In our experimental conditions, the cell survival reached its maximum when the delay between bunches was about 25s. This result explain why no differences in cell survival was found between LDP and conventional beams in our previous study, with a repetition rate of about 1 shot every 30 seconds^5^, leading to an average dose rate in the range of 1.4 Gy/min, roughly as much as conventional beams. This value probably relies on the intrinsic susceptibility of HCT116 cells and may be different with different cell types. Indeed, it has been shown that the W-shaped survival curve profile varies from a cell line to another, with the “peak” occurring between 3 and 60 seconds^14, 15^. We did not observed a W-shaped curve but a bell-shaped curve when varying the pace of bunches. We cannot exclude that with higher repetition rate, cell survival could increase again. This condition will be explored in a next future, when the LDP generation will be upgraded to reach a repetition rate higher than 0.5 Hz. It could also come from the fact that we applied sequences of five LDP bunches on HCT116 cells whereas WE was observed with two bunches. Either the application of more than two short pulses of ionizing radiations could result in the modification of the W-curve profile or it could induce some kind of resonance of the biological mechanisms involved, resulting in a kind of bell-shaped profile. These results suggests that fast fractionation of the dose has an additional impact on the resulting cell survival at constant dose, whose effect varies depending on the numbers and the repetition rate of pulses. The W-effect is the consequence of a particular fast dose fractionation condition, the application of two pulses, whereas the application of several pulses leads to non-monotonic cell survival variation.

Finally, we observed the lowest cell survival for a bunch delay of about 2-3 seconds. Interestingly, at this repetition rate, the cell survival of the radioresistant HCT116 p53^−/−^ cells converged to that of HCT116 WT cells, which was similar to the case when Olaparib was added. This result suggests that, depending on the bunch repetition rate, cell killing obtained with laser pulsed particles could be as efficient as combined therapy using PARP inhibitors. This particular point highlights a therapeutic advantage that laser plasma accelerators could provide compared to continuous therapeutic beams. Indeed, the use of PARP inhibitors in clinical protocols leads, as any drug, to side effects and drug resistance^31, 32^ and drug resistance. Altogether, these results supported the great therapeutic potential of laser plasma technology and pulsed beams of ionizing radiation to provide innovations in radiotherapy for cancer treatment.

## Methods

### Cell culture and chemical treatments

The human colorectal cancer HCT116 WT and p53-/-cells were cultured in McCoy’s5A (Modified) Medium. The human glioblastoma cell lines, SF763 and U87-MG were cultured in Dulbecco’s modified Eagle’s minimum medium with Glutamax (Thermo Fisher Scientific). Cells were grown as monolayers in their respective medium, supplemented with 10% fetal calf serum (PAA) and 1% penicillin and streptomycin (ThermoFisher Scientific) in plastic tissue culture disposable flasks (TPP) at 37°C in a humidified atmosphere of 5% CO_2_ in air. For PARP1 inhibition, cells were incubated one hour in medium supplemented with Olaparib (AstraZeneca, 20 µM diluted in DMSO) to a final concentration of 200 nM before irradiation or oxidative treatment. Untreated cells were exposed to the same volume of DMSO. For hydrogen peroxide (H_2_O_2_) treatment, H_2_O_2_ (Sigma Aldrich 216763, 100 mM in water) was added to culture medium to a final concentration of 1 mM and cells were left exposed for a 10 min incubation time at room temperature.

### Conventional irradiation conditions

X-ray irradiations were performed using a Varian NDI 226 X-ray tube applying as previously described^33^ a tension of 200 kV and an intensity of 15 mA with a dose rate of 1,23 Gy/min. Irradiation with conventional accelerated protons (CAP) were performed using the proton beam at the Curie Institute Proton Therapy Center^34^: an IBA C230 isochronous cyclotron, which generates a proton beam with an initial energy of 235 MeV. The beam energy is lowered to 201 MeV right at the cyclotron output and further reduced down to 20 MeV by sequential polycarbonate and Plexiglas attenuators.

### Laser-driven proton source

The LDP irradiations were performed with laser-driven, TNSA-based proton beam. In a TNSA (Target Normal Sheath Acceleration) proton source, protons are accelerated by the charge separation that is produced at the frontier of an expanding plasma. Such a plasma is created and heated during the interaction between an intense, ultra-short laser pulse and a solid target, where the laser field at high intensity efficiently heats electrons in the target material and increases the accelerating field, hence the proton final energy^21^.

In our experiments, the 200 TW laser source of SAPHIR (Amplitude Technologies) at the Laboratoire d’Optique Appliquée (LOA, Palaiseau, France) was used as a main driver. The laser pulse duration is 26 fs and the laser energy on target is 3 J per shot, for an on-target intensity approaching 10^20^ W/cm^2^. The laser pulse was focused on 5 μm thick Titanium targets, which provided the bulk material for the production of a plasma and the surface where hydrogen ions sit before acceleration. A target is destroyed at each shot. A semi-automatic alignment procedure ensures the alignment of a fresh target after each shot within tolerances, in order to ensure comparable acceleration conditions during the multi-shot irradiation procedure. The initial laser repetition rate was thus lowered from native 5 Hz to single shot, in order to match the pace of the target change, which can be slower or equal to 0.5 Hz (2 seconds between two consecutive shots).

The particle beam shaping, selection and transport was ensured by a set of four permanent magnet quadrupoles^35^. TNSA-based sources produce a multi-specie, highly divergent, laminar and chromatic particle beam. The transport line has been designed to reject electrons, unwanted ion species and to focus the proton emission to a surface of 2 cm^2^ where the biological samples were positioned, 97 cm far from the plasma point.

The quality of the beam during target irradiation (shot-to-shot stability and total deposited dose) was assessed by real-time dosimetry measurements. These are obtained through a transmission ionization chamber (PTW 786, 100 μm water equivalent thickness); the ionization chamber combined with an electrometer (PTW, UNIDOS^®^ E) was previously calibrated with the synchrotron proton beam at the Orsay Protontherapy center (CPO).

Dosimetry measurements in the conditions the present study showed a total dose-per-shot of 0.7 Gy within the 2 cm^2^ irradiation surface. The dose map per unit surface has a uniformity of 20% rms. Laser-accelerated particle bunches hold the temporal signature of the fast accelerating process, which takes place during a time shorter than 1 ps. In the case of MeV protons, the continuous spectrum produces a stretch of the particle bunch during its propagation. Considering the distance between the proton emission point and irradiation point, as well as the available spectrum, total dose in a single shot was delivered at the biological target during τ = 4.8 ns. This corresponded, for a measured dose of 0.7 Gy/shot, to a peak dose rate in the order of 1.5×10^8^ Gy/s^5^.

### Cell survival assay

The cell containers used for irradiation were lumox^®^ dish 35 (SARSTEDT) exhibiting a 25 µm thick lumox^®^ bottom face. Depending on beam transverse profile and position observed on a sensitive film (Image plate, Fujifilm) a 18 mm × 0.9 mm area was delimited on the internal face of the lumox^®^ membrane, where 6×10^4^ cells were seeded. Cells were let grow overnight in 200µl of medium. To generate dose-response survival curves, HCT116 cells were exposed to either 1, 2, 4, 6 or 8 consecutive bunches of LDP or to 1, 2, 3, 4 or 6 Gy of CAP or X-rays, whereas SF763 and U87-MG cell lines were subjected to 2, 5, 7, 10 or 12 consecutive bunches of LDP or 2, 4, 6, 8 or 10 Gy of CAP or X-rays. To generate survival profiles in function of the repetition rate of LDP bunches, HCT116 cells were exposed to series of 5 LDP bunches and glioblastoma cells, SF763 and U87-MG, to series of 9 bunches. After exposure to ionizing radiations, cells were incubated for 3 hours in standard conditions. Cells were harvested with Accutase (Merck), dispatched into 3 different wells of 12-well plates (TPP) in 2.5 ml of medium and grown for five generations corresponding to 5 days for HCT116 and SF763 cells, and 6 days for U87-MG cell line. Cells were harvested with Accutase which then be inactivated with the same volume of 1X PBS (ThermoFisher Scientific) supplemented with 10% fetal calf serum. The final volume was adjusted to 1 ml with 1X PBS and 200µl of each well were dispatched into a non-sterile U-bottom 96-well plate (TPP). In each well, 2 µl of a propidium iodide solution (Sigma, 100 µg/ml in 1X PBS) were added just before flow cytometry counting. Cell acquisition and data analysis were performed using Guava^®^ and GuavaSoft (Merck).

### DNA damage foci immunofluorescent staining

For DNA damage foci analysis, 4×10^5^ cells were seeded on the entire lumox^®^ surface of the dish for LDP irradiation or into Slide Flasks (NUNC) for CAP or X-rays irradiations. Cells were subjected to 3 or 6 consecutive bunches of LDP or to 2 or 4 Gy of CAP or X-rays. The cells were incubated one hour post irradiation at 37°C, then washed with 1X PBS and fixed in 4 % Formalin solution (Sigma) for 15 min at room temperature. The cells were then washed twice with 1X PBS before incubation for 10 min in 1 ml of permeabilization buffer (0.5 % Triton X-100 in 1X PBS). After a 20-min saturation in 1 ml of 1X PBS supplemented with 2% SVF, saturation buffer was entirely removed. Opened Lumox^®^ dishes were placed 5 min under chemical hood to eliminated liquid excess. The Immunostaining surface was delimitated using Dako Pen (Agilent). The cells were incubated for one hour at room temperature with a mouse monoclonal antibody against γH2AX (Merck) diluted in PBS/SVF (1/1000, 30 µl per area). The cells were then washed twice with PBS/SVF before incubation in a 50 µl dilution of secondary antibody (Alexafluor 488 goat anti-mouse antibody, ThermoFisher Scientific, 1/500 in PBS/SVF). After a wash with 1X PBS, the cells were incubated in 1X PBS supplemented with DAPI (ThermoFisher Scientific, 0.1µg/ml) for 10 min at room temperature and were finally washed once again with 1X PBS. Liquid excess was evaporated 5 min under chemical hood. The irradiated Lumox^®^ area was cut following Dako Pen delimitation and was mounted on glass microscopy coverslips. Coverslips were then mounted on slides from Slide Flasks using Prolong^®^ Gold Antifade (ThermoFisher Scientific). Picture Acquisition was performed using DMi8 microscope and LASX software (Leica) at 40x magnification.

### Automated counting of DNA damage foci

Foci were automatically counted in each nucleus, using an ImageJ macro as previously described^33^. Nuclear images were process to correct eventual uneven illumination by dividing the original nuclear image by a duplicate convolved by a gaussian filter. The resulting image intensities were enhanced and smoothed using successively a gamma transform and a median filter. Automated threshold cut was applied, followed by morpho-mathematics operations (close, dilate, fill holes and watershed). The "analyze particle" function was used to isolate individual nuclei and retrieve their contours. The outline of each nucleus was drawn onto the image of DNA repair foci. Foci were characterized as local maxima, and the corresponding ImageJ function was used to identify and count them.

### Protein extraction and Western Blotting

Cells were exposed to Olaparib and/or to H_2_O_2_, medium was removed and cells were harvested and lysed by adding SDS-Urea buffer (62.5 mM TrisHCl pH6.8, 6 mM Urea, 10% Glycerol and 2% SDS) on ice. Cell extracts were sonicated 10 s and protein concentration determined using DC^TM^ Protein Assay (Bio-Rad). Ten micrograms of proteins were loaded and run on NuPAGE 3-8% Tris-Acetate (ThermoFisher Scientific) before being transferred onto 0.45 µm PVDF membrane (Merck). Immunostaining was performed by incubating membrane 1h at room temperature with 1/1000 diluted of rabbit polyclonal anti-Poly-ADP Ribose (TREVIGEN 4336-BPC-100), rabbit monoclonal anti-PARP1 (Cell Signaling Technology CST#9532) or mouse monoclonal anti-GAPDH (Merck, MAB374) antibodies. Goat anti-rabbit and goat anti-mouse IgG antiserums conjugated to peroxidase (SouthernBiotech, 4050-05 and 1031-05 respectively, 1/5000) to reveal protein signals. Images were acquired using G:BOX (Syngene).

### Data analysis

Data analyses were performed using the GraphPad Prism software. The linear quadratic model was applied to fit the survival curves. Depending on experimental design, ratio paired t-test or the two-way ANOVA Tukey’s multiple comparisons tests, with α=0.05, were applied: “*” corresponds to p<0.05, “**” to p<0.001, “***” to p<0.001 and “****” to p<0.0001.

## Acknowledgement

The authors acknowledge the support of OSEO Project No. I0901001W-SAPHIR and of the European Research Council through the X-Five ERC project (Contract No. 339128). F.M.C. aknowledges funding from “France Hadrons” for the CAP irradiation facilities. E. B. acknowledges support from the European Union’s Horizon 2020 research and innovation program (H2020-INFRAIA-2014-2015) under Grant Agreement No. 654148 Laserlab-Europe. This work was also supported by Paris-Saclay University IRS financing, NanoTherad project and EDF Grant No. RB 2017-21 and RB 2018-15.

## Author Contributions

The study was planned and supervised by E.B., A.F., F.M.C., E.D. and V.M. Experimental setup and data acquisition was performed by E.B., O.D., L.P., D.L. and M.C. with help by F.M.C. and A.F. Data evaluation and writing of the publication manuscript was jointly done by E.B., F.M.C. and A.F. with help by E.D. and V.M.

## References

1 Doyen, J., Falk, A. T., Floquet, V., Herault, J. & Hannoun-Levi, J. M. Proton beams in cancer treatments: Clinical outcomes and dosimetric comparisons with photon therapy. Cancer treatment reviews 43, 104–112, doi:10.1016/j.ctrv.2015.12.007 (2016).

2 Durante, M. & Loeffler, J. S. Charged particles in radiation oncology. Nature reviews. Clinical oncology 7, 37–43, doi:10.1038/nrclinonc.2009.183 (2010).

3 Bin, J. et al. A laser-driven nanosecond proton source for radiobiological studies. Applied Physics Letters 101, 243701, doi:10.1063/1.4769372 (2012).

4 Doria, D. et al. Biological effectiveness on live cells of laser driven protons at dose rates exceeding 109 Gy/s. AIP Advances 2, 011209, doi:10.1063/1.3699063 (2012).

5 Pommarel, L. et al. Spectral and spatial shaping of a laser-produced ion beam for radiation-biology experiments. Physical Review Accelerators and Beams 20, 032801 (2017).

6 Tommasino, F. & Durante, M. Proton Radiobiology. Cancers 7, 353 (2015).

7 Fiorini, F. et al. Dosimetry and spectral analysis of a radiobiological experiment using laser-driven proton beams. Physics in Medicine & Biology 56, 6969 (2011).

8 Kraft, S. D. et al. Dose-dependent biological damage of tumour cells by laser-accelerated proton beams. New Journal of Physics 12, doi:DOI:101088/1367-2630/12/8/085003 (2010).

9 Manti, L. et al. The radiobiology of laser-driven particle beams: focus on sub-lethal responses of normal human cells. Journal of Instrumentation 12, C03084 (2017).

10 Raschke, S. et al. Ultra-short laser-accelerated proton pulses have similar DNA-damaging effectiveness but produce less immediate nitroxidative stress than conventional proton beams. Scientific reports 6, 32441, doi:10.1038/srep32441 (2016).

11 Yogo, A. et al. Measurement of relative biological effectiveness of protons in human cancer cells using a laser-driven quasimonoenergetic proton beamline. Applied Physics Letters 98, 053701, doi:10.1063/1.3551623 (2011).

12 Yogo, A. et al. Application of laser-accelerated protons to the demonstration of DNA double-strand breaks in human cancer cells. Applied Physics Letters 94, 181502, doi:10.1063/1.3126452 (2009).

13 Zeil, K. et al. Dose-controlled irradiation of cancer cells with laser-accelerated proton pulses. Applied Physics B 110, 437–444, doi:10.1007/s00340-012-5275-3 (2013).

14 Ponette, V. et al. Hyperfast, early cell response to ionizing radiation. International journal of radiation biology 76, 1233–1243 (2000).

15 Fernet, M. et al. Poly(ADP-ribose) polymerase, a major determinant of early cell response to ionizing radiation. International journal of radiation biology 76, 1621–1629 (2000).

16 Langelier, M. F. & Pascal, J. M. PARP-1 mechanism for coupling DNA damage detection to poly(ADP-ribose) synthesis. Current opinion in structural biology 23, 134–143, doi:10.1016/j.sbi.2013.01.003 (2013).

17 Chen, A. PARP inhibitors: its role in treatment of cancer. Chinese journal of cancer 30, 463–471, doi:10.5732/cjc.011.10111 (2011).

18 de Murcia, J. M. et al. Requirement of poly(ADP-ribose) polymerase in recovery from DNA damage in mice and inLcells. Proceedings of the National Academy of Sciences 94, 7303–7307, doi:10.1073/pnas.94.14.7303 (1997).

19 Nile, D. L., Rae, C., Hyndman, I. J., Gaze, M. N. & Mairs, R. J. An evaluation in vitro of PARP-1 inhibitors, rucaparib and olaparib, as radiosensitisers for the treatment of neuroblastoma. BMC cancer 16, 621, doi:10.1186/s12885-016-2656-8 (2016).

20 Ray Chaudhuri, A. & Nussenzweig, A. The multifaceted roles of PARP1 in DNA repair and chromatin remodelling. Nature reviews. Molecular cell biology 18, 610–621, doi:10.1038/nrm.2017.53 (2017).

21 Wilks, S. C. et al. Energetic proton generation in ultra-intense laser–solid interactions. Physics of Plasmas 8, 542–549, doi:10.1063/1.1333697 (2001).

22 Bulanov, S. V. & Khoroshkov, V. S. Feasibility of using laser ion accelerators in proton therapy. Plasma Physics Reports 28, 453–456, doi:10.1134/1.1478534 (2002).

23 Malka, V. et al. Practicability of protontherapy using compact laser systems. Medical Physics 31, 1587–1592, doi:doi: 10.1118/1.1747751 (2004).

24 Masood, U. et al. A light-weight compact proton gantry design with a novel dose delivery system for broad-energetic laser-accelerated beams. Physics in medicine and biology 62, 5531–5555, doi:10.1088/1361-6560/aa7124 (2017).

25 Ponette, V. et al. Pulse exposure to ionizing radiation elicits rapid changes in cellular radiosensitivity. J Comptes rendus de l’Academie des sciences. Serie III, Sciences de la vie 319 6, 505–509 (1996).

26 Wei, H. & Yu, X. Functions of PARylation in DNA Damage Repair Pathways. Genomics, Proteomics & Bioinformatics 14, 131–139, doi:https://doi.org/10.1016/j.gpb.2016.05.001 (2016).

27 <https://clinicaltrials.gov>(

28 Dungey, F. A., Löser, D. A. & Chalmers, A. J. Replication-Dependent Radiosensitization of Human Glioma Cells by Inhibition of Poly(ADP-Ribose) Polymerase: Mechanisms and Therapeutic Potential. International Journal of Radiation Oncology • Biology • Physics 72, 1188–1197, doi:10.1016/j.ijrobp.2008.07.031 (2008).

29 Shervington, A. et al. Telomerase subunits expression variation between biopsy samples and cell lines derived from malignant glioma. Brain Research 1134, 45–52, doi:https://doi.org/10.1016/j.brainres.2006.11.093 (2007).

30 Makvandi, M. et al. A PET imaging agent for evaluating PARP-1 expression in ovarian cancer. The Journal of Clinical Investigation 128, 2116–2126, doi:10.1172/JCI97992 (2018).

31 Bitler, B. G., Watson, Z. L., Wheeler, L. J. & Behbakht, K. PARP inhibitors: Clinical utility and possibilities of overcoming resistance. Gynecologic Oncology 147, 695–704, doi:https://doi.org/10.1016/j.ygyno.2017.10.003 (2017).

32 Zhou, J. X., Feng, L. J. & Zhang, X. Risk of severe hematologic toxicities in cancer patients treated with PARP inhibitors: a meta-analysis of randomized controlled trials. Drug design, development and therapy 11, 3009–3017, doi:10.2147/DDDT.S147726 (2017).

33 Bayart, E. et al. Enhancement of IUdR Radiosensitization by Low-Energy Photons Results from Increased and Persistent DNA Damage. PloS one 12, e0168395, doi:10.1371/journal.pone.0168395 (2017).

34 Calugaru, V. et al. Radiobiological Characterization of Two Therapeutic Proton Beams With Different Initial Energy Spectra Used at the Institut Curie Proton Therapy Center in Orsay. International Journal of Radiation Oncology • Biology • Physics 81, 1136–1143, doi:10.1016/j.ijrobp.2010.09.003 (2011).

35 Schillaci, F. et al. Characterization of the ELIMED Permanent Magnets Quadrupole system prototype with laser-driven proton beams. Vol. 11 (2016).

